# Context-Dependent Genetic Regulation

**DOI:** 10.1101/173906

**Authors:** Kyung Hyuk Kim, Venkata Siddartha Yerramilli, Kiri Choi, Herbert M. Sauro

## Abstract

Cells process extra-cellular signals with multiple layers of complex biological networks. Due to the stochastic nature of the networks, the signals become significantly noisy within the cells and in addition, due to the nonlinear nature of the networks, the signals become distorted, shifted, and (de-)amplified. Such nonlinear signal processing can lead to non-trivial cellular phenotypes such as cell cycles, differentiation, cell-to-cell communication, and homeostasis. These nonlinear phenotypes, when observed at the cell population levels, can be quite different from the single-cell level observation. As one of the underlying mechanisms behind this difference, we report the interplay between nonlinearity and stochasticity in genetic regulation. Here we show that nonlinear genetic regulation, characterized at the cellular population level, can be affected by cell-to-cell variability in the regulatory factor concentrations. The observed genetic regulation at the cell population is shown to be significantly dependent on the upstream DNA sequences of the regulator, in particular, 5’ untranslated region. This indicates that genetic regulation observed at the cell population level can be significantly dependent on its genetic context, and that its characterization needs a careful attention on noise propagation.

**One Sentence Summary:** Genetic regulation observed at the cell population level can be significantly affected by cell-to-cell variability in the regulatory factor copy numbers, indicating that the observed regulation is dependent on 5’ UTR of the regulator coding gene.

## 1 Introduction

Genetic regulation often shows highly nonlinear patterns, for example, on-off switching with respect to the concentration levels of regulatory factors (transcription factors) (*1*). In addition, the factor concentration levels fluctuate significantly due to intracellular random biological reactions (*2*). This randomness – stochasticity – and the nonlinearity can provide a variety of noise-related cellular phenotypes (*3–6*). Here, we investigate the interplay between stochasticity and nonlinearity. We report that genetic regulation that are measured at the cell population level and at the single cell level can be significantly different because of the interplay (*5, 7*). Considering that stochasticity observed in a transcription factor concentration level can be controlled without altering a pair of the transcription factor and its specific promoter (*8–12*), our result implies that the genetic regulation is dependent on genetic context.

Consider the nonlinear signal process shown in Fig. 1A. When the process is linear, the mean values of the input and output are located on top of the response curve. However, when the process is nonlinear, for example, in the curve-down response, the output signal mean value is placed lower than the response curve (Fig. 1A, middle). This is due to the fact that the output signals are distorted by the nonlinear signal processing (Fig. 1B).

This nonlinear effect can be understood mathematically with the Jensen’s inequality: For example, the graph of a convex function lies below all its secant lines (Fig. 1C):

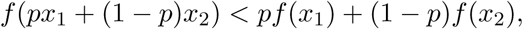

for 0 < *p* < 1 and when *x*_1_ and *x*_2_ get closer to each other, the inequality becomes weaker. This inequality statement can be re-phrased by considering *x* as a random variable. The value of a convex function at the mean value of *x* is smaller than the mean value of the function:

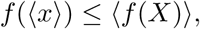

where 〈∙〉 represents the mean value. When the value of *x* is less noisy (smaller randomness), the inequality becomes weaker. This indicates that the output mean value can be dependent on the strength of the input signal noise. Thus, when averaging cellular signals over the cell population, the population responses can be significantly different from the individual cellular responses due to interplay between stochasticity and nonlinearity.

**Figure 1:**
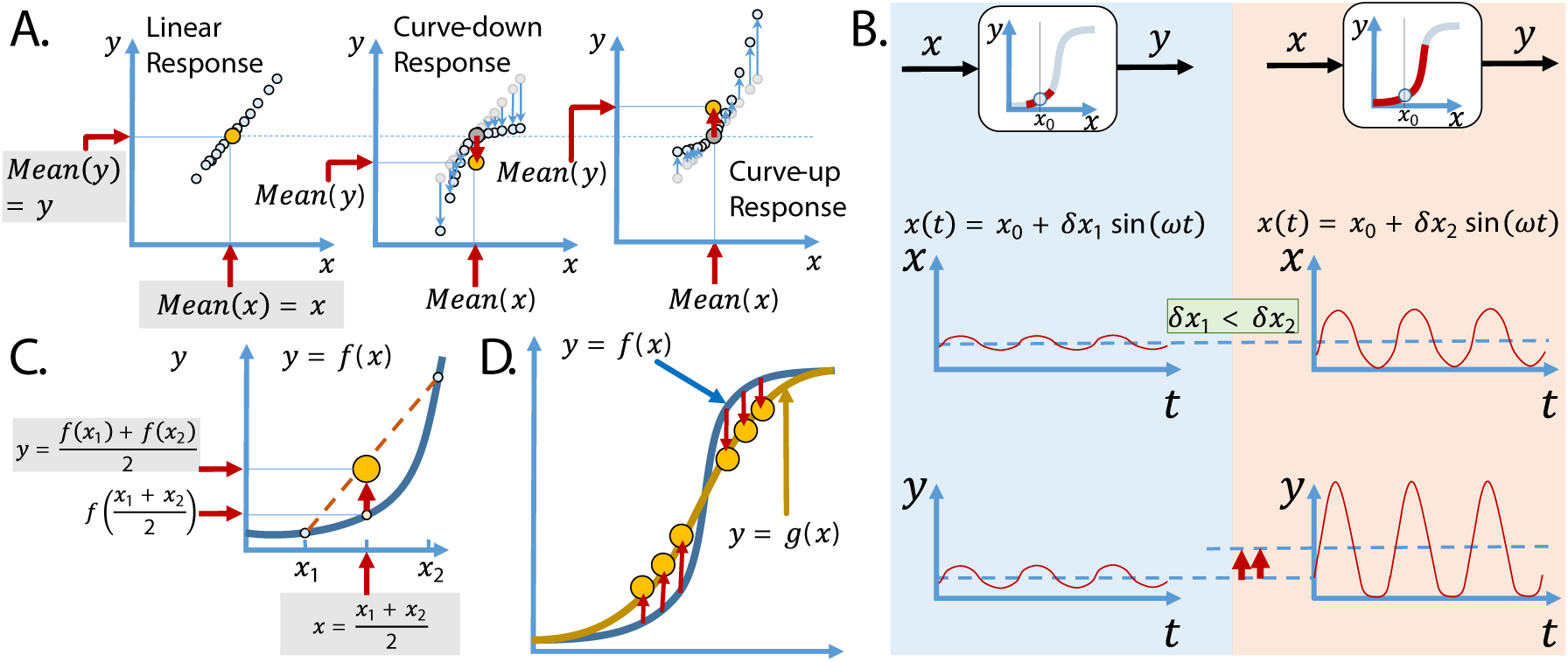
Nonlinear signal processing: (A) The mean value of an output signal is affected by the shape of the input-output response curve (i/o curve). The light blue circles correspond to single observations and the orange circles to the mean values of all the observations. When the i/o curve is linear, the output mean value is placed on top of the response curve (here, straight line) (when neglecting noise within the signal processing). When the i/o curve is nonlinear, the output mean value is placed below or above the response curve depending on its curvature. (B) Input signals to a nonlinear processor are sinusoidal and its output signals are distorted when the amplitude of the input oscillation is large enough to see the nonlinearity. (C and D) Jensen’s inequality shows the averaged input-output response dependent on the input signal noise strength.

Consider sigmoidal input-output response as shown in Fig. 1D. For the convex (curve-up) region of the response, the output mean value becomes larger than the curve itself and opposite for the concave region. This indicates that the response curve measured in the average, such as at the cell culture (population) level, can be significantly different from the response measured at the single cell level. Here we will investigate such differences in *E. coli* genetic regulation.

Genetic regulation by transcription factors (TFs) has been characterized typically based on cellular culture at the population level, for example, by using a microplate photometer (*13*). TF regulation can be highly nonlinear and the intracellular TF copy number can show significant cell-to-cell variability. Here, we will consider the genetic regulation of heterogenous LuxR activator in *E. coli* observed at the population and single cell levels. We show that the regulation patterns observed at both the levels are different, implying that the genetic regulation measured at the population level can be context-dependent, i.e., dependent not only on the pair of LuxR and its specific promoter (*plux*) but also on other genetic components such as the 5’ UTR and promoter region of the *luxR* coding gene. Thus, genetic regulation needs to be carefully characterized by considering the strength of the gene expression noise and its effect on the regulation pattern.

## 2 Results

### 2.1 Orthogonal control of noise and mean levels

To observe the interplay between nonlinearity and stochasticity in relation to genetic regulation, it is important to have an experimental method to control the noise strength in intracellular TF concentrations. Such noise control has been performed in various types of cells (*8–12*). Our previous study (*14*) adopted the dual control of transcription and translation efficiencies to achieve orthogonal control of mean (*m*) and noise levels (*n*). The noise level – defined here by coefficient of variation, i.e., standard deviation divided by mean – was shown to be varied by a factor of ~3 with the p15A origin of replication (ORI) (Fig. 2B, KK16A strain series) (*14*), where the histograms of GFP concentrations were shown to follow the Gamma distribution and the relationship between *m* and *n* satisfied

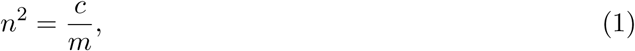

with *c* a constant (Fano factor) while varying the IPTG concentrations for a given ribosome binding site (RBS; BBa_B0034), and with *c* decreasing while adding longer adenine spacers downstream of a given RBS (BBa_B0034) (*14*).

**Figure 2:**
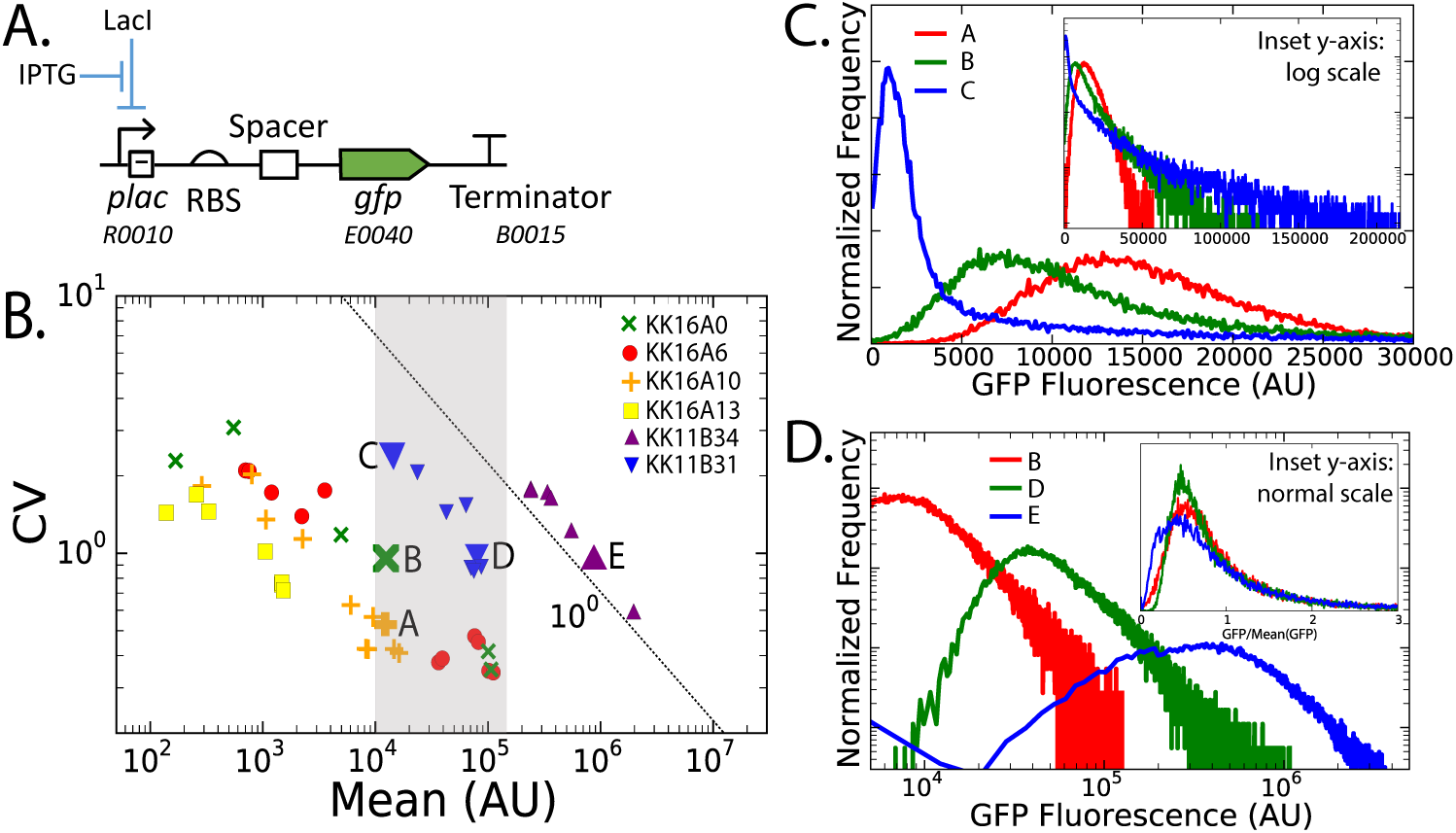
Orthogonal control of gene expression noise and mean values: (A) The *gfp* expression cassette is integrated in plasmids (pGA3K3 ORI: p15A and pGA1A3 ORI: pMB1). The transcription and translation efficiencies were controlled by varying IPTG concentrations and either by using different RBSs or adding spacers between a given RBS (B0034) and *gfp*. The SBOL compliant figure is shown (*27*). (B) The noise strength of the *gfp* expression and its mean value were plotted for different control methods. KK16A strain series have pGA3K3-backbone plasmids, and different spacers (no spacer and 6, 10, and 13 repeats of adenine) were used downstream of RBS B0034. KK11B strain series have pGA1A3 backbone plasmids, and different RBSs (B0034 and B0031) were used without any adenine repeat spacer. The data points A, B, and C correspond to orthogonal control of the *gfp* expression noise level (C), and the data points B, D, and E to orthogonal control of the expression mean level (D).

Here, we achieved a further increase in *n* without changing *m* by using different ORIs. Fig. 2B (gray region) shows that when using the ORI of the higher copy number plasmid, pMB1, *n* increases another ~ 2 fold change. As a net, we have achieved ~ 6 fold increase in *n* without changing *m* (Fig. 2B) via triplet control of transcription and translation efficiencies and plasmid copy number. Fig. 2B shows that the graph of *m* vs. *n* tends to shift to the right when replacing p15A ORI to pMB1 ORI as well as when using stronger RBSs. This observed shift indicates that the intrinsic gene expression processes – here the *plac* regulated transcription-translation processes – are much stronger than the extrinsic noise that is mostly originated from cell doubling (*15*). If the plasmid expression were not used but instead genomic expression investigated, the intrinsic noise would have been buried by the extrinsic noise as observed for typical *E. coli* transcription factors with their copy numbers higher than ~ 10 (*15, 16*) (confer to the study on yeast transcriptome (*17*), where intrinsic noise is strong enough to be observed for most transcription factors because the extrinsic noise level is lower than *E. coli* due to its longer doubling time). The minimum noise level that we observed in the GFP signals is slightly larger than 0.1, which is the minimal noise level where the extrinsic noise becomes dominant in *E. coli* (*15*).

The value of *n* is significantly dependent on the tail distributions of gene expression level histograms. Fig. 2C (inset plot) shows the case of similar *m* but different *n* values, where the tail distributions become longer as *n* increases. Fig. 2D shows the case of different *m* but similar *n* values, where the tail distributions are very similar. This implies that the tail distributions follow the similar probability distribution function for the Points B, D, and E and they are a dominant factor determining the noise strength.

### 2.2 Noise-dependent genetic regulation

To observe the effect of cell-to-cell variability in TF copy numbers on the downstream regulation pattern, the LuxR::Venus fusion as shown in Fig. 4A was used to monitor the copy number variation of LuxR regulators. To maximize the effect of the cell-to-cell variability (noise) onto the downstream regulation, we used a strong ribosome binding site, B0034, for LuxR expression, and high copy number plasmids by using the pMB1 origin (pGA1A3).

We suspect that a time-delay in activating mCherry expression and maturating the fluorescent proteins can de-synchronize the two signals of Venus and mCherry, making extremely difficult to observe the LuxR-activation pattern. Single-cell microscopy experiments were performed (refer to the Materials and Methods) and confirmed the time-delay (Fig. 3A,D). When the time delay is taken care of (refer to the Materials and Methods), the nonlinear regulation transfer curve was observed (Fig. 3C,F). The obtained transfer curve was, however, significantly noisy, because of the sample size limit of approximately 100 lineage trajectories in the microscopy experiments.

**Figure 3:**
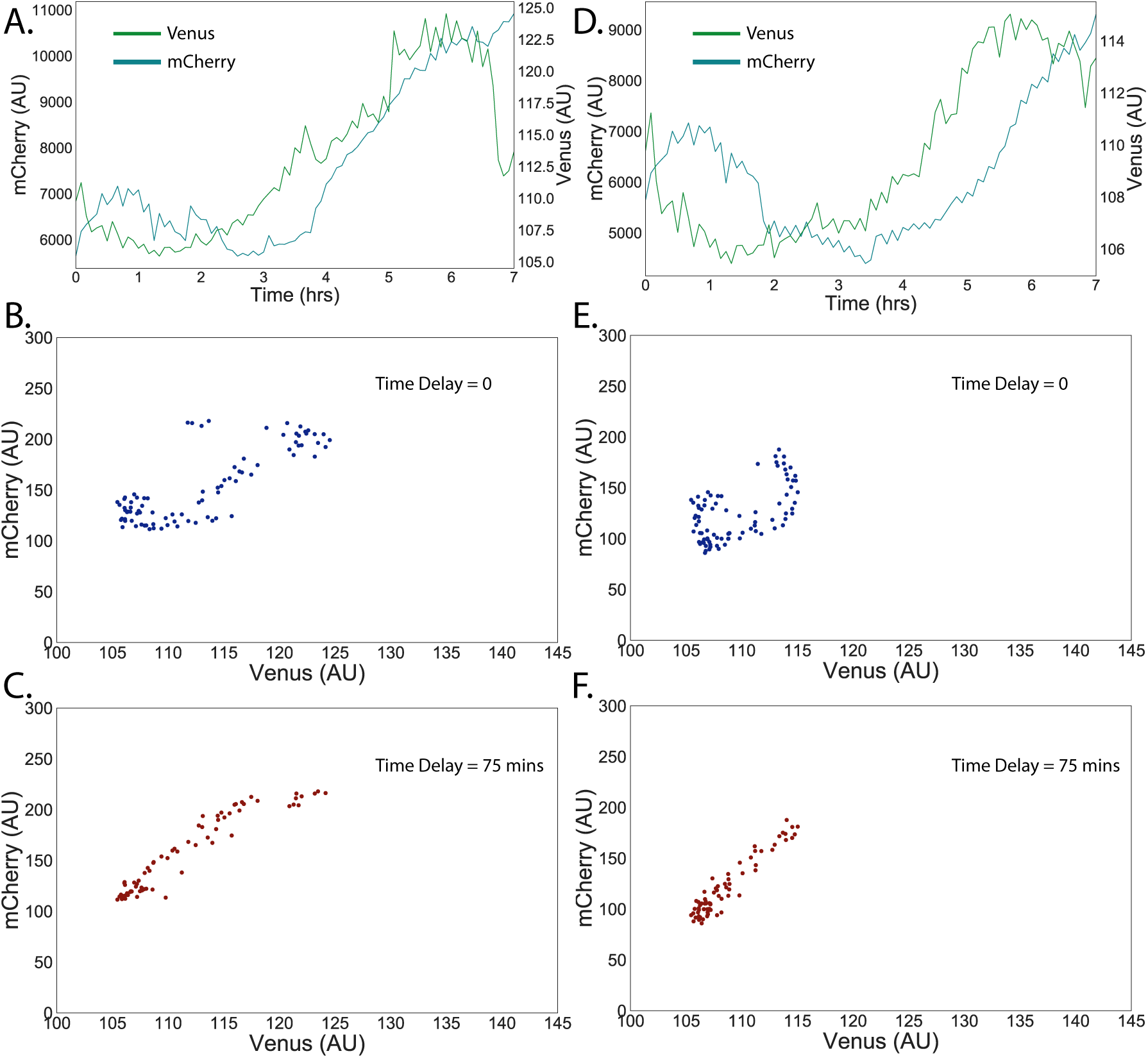
Time delay in LuxR regulation: Two cell lineages, (A, B, C) and (D, E, F), are shown. (A,D) The blue line represents mCherry and the green line Venus. The fluorescence signals were measured for ~7 hours with 5 min interval, corresponding to ~6 cell divisions. The time delay of ~ 75 mins in the activation (B,E) and the temporal change in mCherry signals (refer to the Materials and Methods) are taken considered and then the regulation pattern (C,F) was recovered.

To overcome these issues of desynchronization and limited sample size, we used overnight cultures and their fluorescence signals were measured via flow cytometry. Based on the data obtained via flow cytometry, we could observe the similar nonlinear regulation pattern as shown in Fig. 4. By using the pMB1 ORI and the strong RBS B0034, we could achieve the cell-to-cell variability of the LuxR copy numbers that is strong enough to cover the nonlinear regulation region (Fig. 4B (red and blue)). The population average (filled circle) was systematically deviated from the single cell trend: At the single cell level, the density plot of venus vs. mCherry shows how probable the genetic regulation is, and thus the ridge line of the density plot indicates the most probable regulation pattern. The population average values for the cases of 7 different IPTG concentrations were deviated from the single-cell level genetic regulation pattern systematically. This confirms that the population-level and single-cell-level experiments can show different genetic regulation patterns via interplay between stochasticity and nonlinearity. In addition, the cell-to-cell variability of LuxR copy numbers can be in principle controlled by changing the *upstream* of the *luxR* gene (RBSs and *lac*-promoter) as well as the plasmid ORI, without altering LuxR-coding gene and its specific promoter (*plux*). This manifests that the LuxR regulation pattern, measured at the population level, is significantly dependent on *genetic context*.

**Figure 4:**
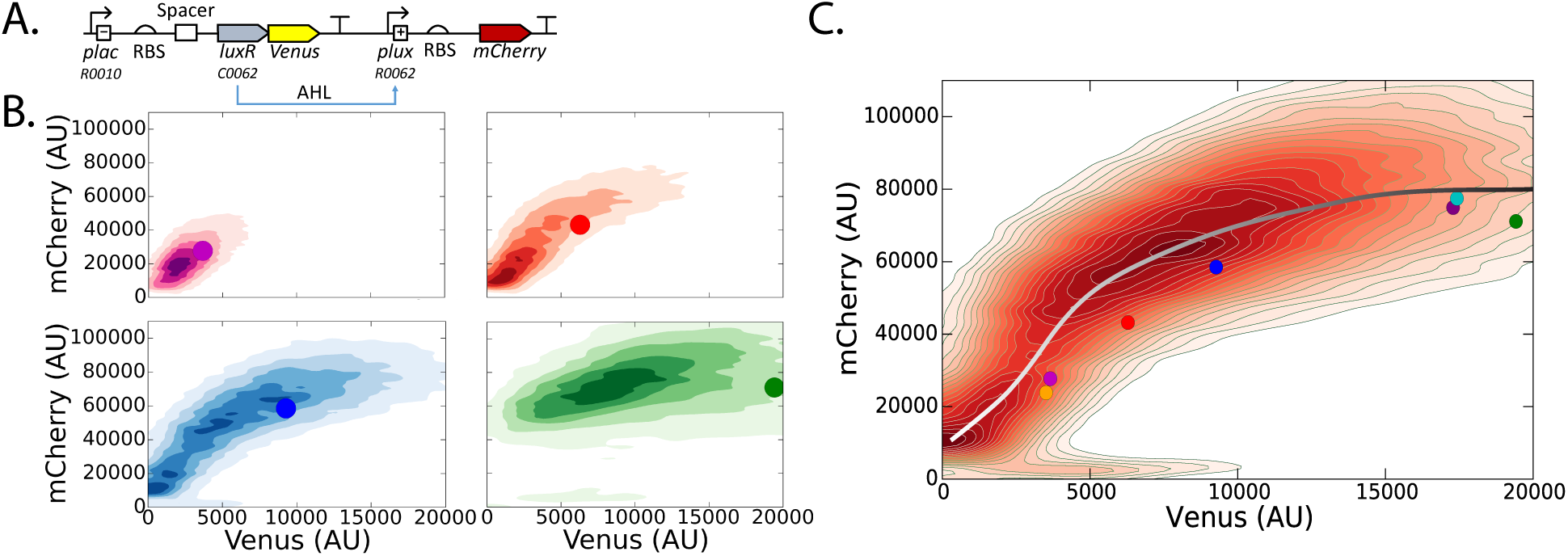
Effect of gene expression noise on genetic regulation: (A) LuxR::Venus fusion activates mCherry expression. RBS is B0034 and the circuit is integrated in pGA1A3 backbone. (B) mCherry and Venus signals were measured via flow cytometry and their density plot were generated for [IPTG] = 0.022, 0.025, 0.063, 0.13 mM (top-left, top-right, bottom-left, bottom-right). (C) Multiple flow cytometry data for 7 different IPTG concentrations were combined. The filled circles are the mean values of Venus and mCherry signals for given [IPTG] (B and C).

## 3 Discussions

### 3.1 Intrinsic vs. extrinsic noise

Our synthetic gene circuits were integrated in plasmids and then *E. coli* was transformed with the plasmids. Fig. 2 shows that the mean and noise levels of the gene expression are inversely related (Eq. 1). This implies that the gene expression from the plasmids does not suppress intrinsic circuit dynamics: In the *E. coli* transcriptome study (*15*), the intrinsic noise was completely suppressed by the extrinsic noise mostly when transcription factor copy numbers were larger than 10 (refer to Fig. 2B in (*15*)). This means that biological processes faster than cell doubling time (extrinsic noise fluctuates typically with the cell doubling time) such as transcription-translation are averaged out. Therefore, to study these processes, it is necessary either to use fast-responsive probes (*18*) or to come up with methods that amplify the intrinsic processes. Encoding genetic systems of interest in plasmids amplifies the signal strength without suppressing the intrinsic processes such as transcription-translation, and this allows us to investigate *E. coli* gene expression with flow cytometry and fluorescence microscopy without resorting to single-molecule fluorescence microscopy (*15, 19*).

### 3.2 Orthogonal noise control

We could achieve ~ 6 fold increase in the noise level without changing its mean level. In principle, the fold increase can be enhanced up to ~ 9 fold. If the gene circuit is placed in a plasmid and/or a strain that can provide tighter control of *plac*, purple triangles shown in Fig. 2B can be extended into the gray-filled area. One possible experiment is that *plac* is replaced to *T7lac* and the circuit is integrated in the pET28a plasmid and Rosetta™ (DE3) *E.coli* is transformed with the plasmid.

### 3.3 Identical tail distributions for the same noise level

Points B, D, and E in Fig. 2B shows an identical tail distribution when rescaled to their mean values equal to one. This scale invariance implies that the tail distributions follow the same functional form. Our previous study showed that the distribution functions for the case of the KK16 series closely follow the Gamma distribution function, 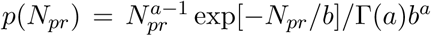, with *b* the translational burst size (*15, 20*). The mean values of GFP was dominantly affected by the use of different RBSs, i.e., different values of *b*. Here, since the KK11 series shows the identical tails to the ones observed with the KK16 series, the tail part of the KK11 series distributions follows the gamma distribution and the translational burst size, *b*, will be the factor that changes the mean values of Points B, D, and E. Considering the fact that the Points E and B correspond to the same RBS (i.e., translation efficiency is identical) and that the burst size *b* – number of proteins synthesized from a single RNA (from its birth to death) – is dependent on both the translation efficiency and RNA lifetime, we can conjecture that RNA lifetime was affected by changing the plasmid replication origin. Based on our mathematical model (Materials and Methods and Supplementary Materials: Mathematica notebook), the plasmid copy number fluctuations did not (if any, negligibly) affect the Fano factor, implying that *b* does not depend on the plasmid copy number fluctuations. Thus, we attribute the observed identical tail distributions to the change in the RNAse activity, which is directly related to the RNA lifetime and thus affects the burst size *b*. Further experiments on RNA stability is suggested by using qRT-PCR to verify this conjecture.

## 4 Materials and Methods

### 4.1 Genetic circuits and strains

Genetic components are mostly BioBrick parts (http://parts.igem.org). Venus and its fusion linker are obtained from (*21*) and mCherry from pNS2-σVL (Addgene Plasmid 26756). The designed gene circuits were constructed via the Gibson assembly method (*22*). All the gene circuits are integrated into either pGA3K3 (p15A origin) and pGA1A3 (pMB1) and *Escherichia coli* MG1655Z1 and NEB Turbo were used as expression and cloning strains, respectively. MG1655Z1 constitutively overexpresses LacI from its chromosome (lacI^*q*^).

### 4.2 Cell Growth, Flow Cytometry and Microscopy Measurements

The *E. coli* strains were grown in 2 mL Luria-Bertani (LB) media (Becton Dickinson) with kanamycin 50 *μ*g/mL at 37°C and 300rpm in a shaker. When OD600 reaches 0.2, the cultures were diluted 1:200 into 200 *μ*L prewarmed fresh M9 media (Teknova 2M1990) in 96-well plates (Costar 3904). The media in each well contains kanamycin 50 *μ*g/mL and different IPTG concentration (0 mM,0.02~1 mM). The M9 cultures were grown to OD600=0.3-0.4 in a shaker (37°C, 300 rpm).

Flow cytometry measurements: A Sony Biotechnology ec800 flow cytometer was used with a 525 nm filter and a 488 nm excitation laser for GFP fluorescence, and a 595 nm filter and a 561 nm excitation laser for mCherry. The cultures grown in M9 were diluted 1:4 in 1xPBS. 100,000 events were collected for each culture and gated by using a 2-D normal distribution by using python package FlowCytometryTool (http://gorelab.bitbucket.org/flowcytometrytools). Well-to-well contamination was prevented by executing the Medium Flush cycle after each sampling. When computing the mean and noise levels of GFP signals of KK16 and KK11 series, background fluorescence was removed by using the mean and noise levels of GFP signals (*14*).

Microscopy Measurements: Cells were inoculated into 2 mL M9 media from -80°C freezer stocks and grown overnight at 37°C and 250 rpm shaking. Overnight cultures were diluted into 1 mL of M9 media supplemented with 0.1 mM IPTG at a 1:1000 ratio and were incubated at 37°C for about 3 hours until OD = 0.05. 2 *μ*L of induced cells were spotted onto an agarose pad on a slide and then covered with a coverslip and sealed using VaLP (1:1:1 vaseline/lanolin/paraffin).The agarose pads were prepared by pouring 1 mL M9 media supplemented with 0.2 % low-melting-point agarose,0.1 mM IPTG and 0.1 mg Carbenicillin into into 2 cm x 2 cm wells cut into a rubber gasket sealed onto a microscope slide (3” x 1”) using the VaLP. Imaging was performed using a Nikon Ti-E inverted wide-field fluorescence microscope with a large-format scientific complementary metaloxide semiconductor camera (NEO, Andor Technology) and controlled by NIS-Elements. Samples were kept at 30 C throughout the imaging process using an environmental chamber. The total imaging time for each experiment was 10 hours during which both bright-field and fluorescent images were captured every 3 minutes.

### 4.3 Image Pre-Processing

Raw microscope image data have been processed through a MATLAB-based software SuperSegger (*23*), a software suite designed for automatic cell segmentation and fluorescence quantification for high-throughput fluorescence microscopy. SuperSegger creates a MATLAB matrix file (.mat) as an output for each image set, which contains various quantities per individual cell per time point. This file was used as an input for our scripts which further processed the data to obtain the results. All of our scripts are based on Python, and utilize NumPy (*24*) for array manipulation, matplotlib (*25*) and seaborn (*26*) for data visualization, and h5py for handling MATLAB file.

### 4.4 Cell Lineage Tracking

The output of SuperSegger contains information on parent and daughter cells for each cell. We used this information to reconstruct a complete map of cell lineage and analyze fluorescence statistics based on that. Prior to running cell lineage tracking, the output of SuperSegger segmentation process has been visually inspected for obvious artifacts (i.e. air bubbles) which are marked to be excluded. Our Python script initially reads all cell data and then purges cells that are marked to be excluded. The tracking process searches for all the daughter cells given the initial parent cells, which is to further reduce potential artifacts. Then, for each lineage, a full time-course measurements of mCherry and Venus fluorescence levels are plotted over the entire duration of microscope measurement in order to provide details on long-term correlation between mCherry and Venus signals. The fluorescence levels of mCherry and Venus are scaled to provide better comparison.

### 4.5 Time Delay Correlation

After obtaining the complete map of cell lineages, we approximated the amount of phase difference between mCherry and Venus signals and generated plots depicting Venus-mCherry response. Because we believed the rate of changes of mCherry level (*d*[*mCherry*]*/dt*) is not negligible, we decided to run 1-dimensional cubic (spline of the third order) interpolation on our mCherry signal time trajectories and obtain their first order derivatives of the interpolation functions. We then plotted out *d*[*mcherry*]*/dt* + *b* [*mcherry*] against Venus signal to obtain corrected response curve, where b is a constant related to cell doubling time. In order to approximate the phase difference between mCherry and Venus signals, we tested 0–75 minute delay and chose the best value for phase difference through visual inspection of the resulting response curves.

### 4.6 Plasmid Copy Number Fluctuations and Their Contribution to the Fano Factor

Consider that the plasmid copy number *N_pl_* follows the distribution *P* (*N_pl_*). The mean protein copy number can be calculated as:

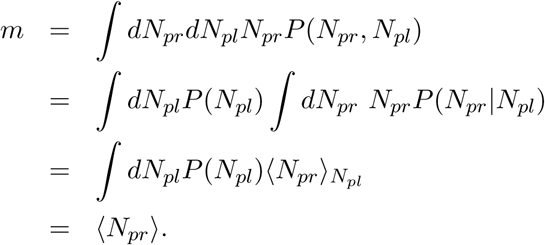

The variance of the protein copy number is expressed as:

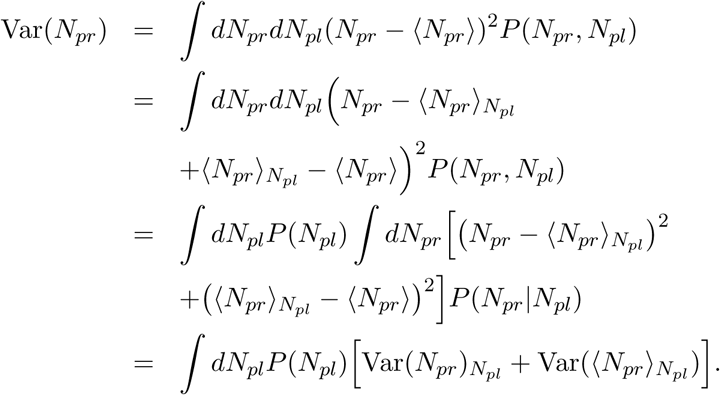

The second term appears because of the fluctuations in the plasmid copy number and if we assume that the mean copy number of proteins is linearly dependent on the copy number of plasmids (i.e., 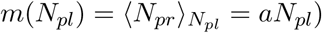, the second term is directly related to the variance of *N_pl_*:

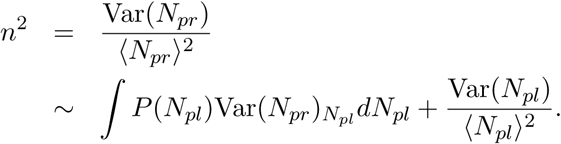

If the plasmid copy number is constant (i.e., no fluctuations), the second term vanishes. The Fano factor could increase due to the plasmid copy number fluctuations. The common sense is that the high copy number plasmids can have smaller copy number fluctuations relative to the mean copy number. Thus, the second term that is related to the plasmid copy number fluctuations needs to be smaller for the case of the high copy number plasmids, which is opposite to the observation. Thus, the effect of the plasmid copy number fluctuations on the changes in the Fano factor values can be minimal.

## 5 Supplementary Materials

Supplementary Notes: DNA sequence information and mathematical models are described. Supplementary Movies: The fluorescent signals of Venus and mCherry emitted from the KK61 strain were taken via microscopy (Materials and Methods). Note that red color corresponds to Venus and yellow color to mCherry.

Supplementary Files: The Mathematica notebook file includes the analysis of the relationship between *n* and *m* based on the mathematical model described in the Supplementary Notes.

## Acknowledgements

We would like to Xiaoliang Sunney Xie and Timothy K Lu for providing plasmids containing Venus and fusion proteins. We would also like to thank Joo-Young Lee for his advice and suggestions. We would also thank Dr. Wiggins and his lab for allowing us to use the microscope.

## 9 Data and Materials Availability

Python script files used to construct cell lineages based on microscopy images can be obtained from https://github.com/kyunghyukkim/microscopy.git.

## Supplementary Notes

### 1 DNA sequences of 5’ UTR regions

Table 1 shows the DNA sequences that were used for the region of ribosome binding sites along with different kinds of spacer sequences:

**Table 1:**
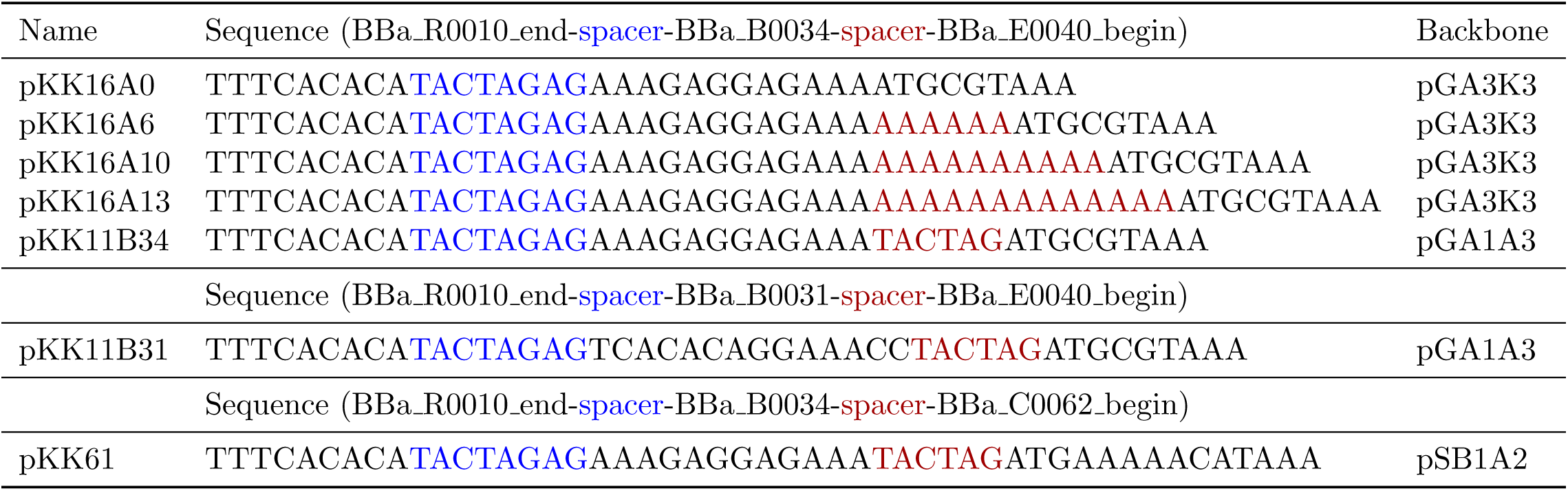
RBS region DNA sequences

### 2 Plasmid information

**Table 2:** RBS region DNA sequences

| Name | Gene Circuit | Backbone |
| --- | --- | --- |
| pKK16(A)<sub>n</sub> | R0010-B0034(A)<sub>n</sub>-E0040-B0015 | pGA3K3 |
| pKK11B34 | R0010-B0034-E0040-B0015 | pGA1A3 |
| pKK11B31 | R0010-B0031-E0040-B0015 | pGA1A3 |
| pKK61 | R0010-B0034-C0062-Venus-B0015-R0062-B0032-mCherry-J61048 | pSB1A2 |

#### 2.1 Mathematical Model

A two-state model for the *lac*-promoter (*1–3*) is considered to describe the promoter active and inactive states. Transcription and translation processes and plasmid copy number fluctuations are considered with simple birth-death processes to show how the fluctuations affect the protein level noise:

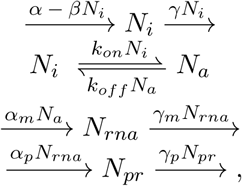

where *N_i_* denotes the number of inactive promoters, *N_a_* that of active promoters, *N_rna_* the RNA copy number, and *N_pr_* the protein copy number. Here, the synthesis rate of the plasmid, i.e., the plasmid replication, is typically dependent on the plasmid copy number; for example, negative feedback has been used to stabilize the plasmid copy number (*4*). Here, we modeled the negative feedback by introducing the term –*βN_i_*, which can be considered as a linearized version of non-linear negative feedback.

This model can be further simplified based on our experimental results. In the main manuscript, we already discussed that the plasmid copy number fluctuations are not the dominant factor for the observed increase in the Fano factor when switching from the medium-low copy number plasmid to the high copy number plasmid. Thus, we will neglect the plasmid copy number fluctuations in the above model and keep the rest of the reactions:

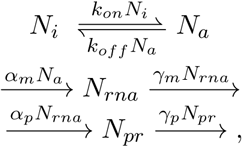

The noise level, *n*, of *N_pr_* was computed numerically from the analytical solutions that were obtained by using the Mathematica (Supplementary File). The relationship between *n* and *m* are investigated by numerically computing these values for different parameter values. Our conclusion was that for the biological reasonable values of the parameters for *E. coli* the change in the Fano factor could be explained by the changes in *γ_m_* and *γ_p_. γ_p_* is typically dependent on cell doubling time because the protein is quite stable comparing to the cell doubling time. Thus, we concluded that *γ_m_* is changed when using the different plasmid backbones, pGA1A3 and pGA3K3.

